# Genome of octoploid plant maca (*Lepidium meyenii*) illuminates genomic basis for high altitude adaptation in the central Andes

**DOI:** 10.1101/017590

**Authors:** Jing Zhang, Yang Tian, Liang Yan, Guanghui Zhang, Xiao Wang, Yan Zeng, Jiajin Zhang, Xiao Ma, Yuntao Tan, Ni Long, Yangzi Wang, Yujin Ma, Yu Xue, Shumei Hao, Shengchao Yang, Wen Wang, Liangsheng Zhang, Yang Dong, Wei Chen, Jun Sheng

## Abstract

Maca (*Lepidium meyenii* Walp, 2n = 8× = 64) of Brassicaceae family is an Andean economic plant cultivated on the 4000-4500 meters central sierra in Peru. Considering the rapid uplift of central Andes occurred 5 to 10 million years ago (Mya), an evolutionary question arises on how plants like maca acquire high altitude adaptation within short geological period. Here, we report the high-quality genome assembly of maca, in which two close-spaced maca-specific whole genome duplications (WGDs, ∼ 6.7 Mya) were identified. Comparative genomics between maca and close-related Brassicaceae species revealed expansions of maca genes and gene families involved in abiotic stress response, hormone signaling pathway and secondary metabolite biosynthesis via WGDs. Retention and subsequent evolution of many duplicated genes may account for the morphological and physiological changes (i.e. small leaf shape and loss of vernalization) in maca for high altitude environment. Additionally, some duplicated maca genes under positive selection were identified with functions in morphological adaptation (i.e. *MYB59*) and development (i.e. *GDPD5* and *HDA9*). Collectively, the octoploid maca genome sheds light on the important roles of WGDs in plant high altitude adaptation in the Andes.

## INTRODUCTION

Maca (*Lepidium meyenii* Walp) is a Brassicaceae crop and herbal plant cultivated in the 4000-4500 meters puna agro-ecological zones in central Andes (http://ecocrop.fao.org/; Supplemental Fig. 1A). Its potential health benefits, particularly to reproduction and fertility, have drawn great investments from pharmacological and nutritional research in recent years (Piacente et al. 2002; Shin et al. 2010). On account of its restricted cultivation area at high altitude, maca manifests robust endurance to the extreme environmental stresses, including cold, strong wind and UV-B exposure. Even though many genes and gene families involved in abiotic stress response have been identified in various plants (Hoffmann and Willi 2008), it is still an open question how plants such as maca adapted to the high altitude Andean environment from an evolutionary genomics standpoint.

**Figure 1.**
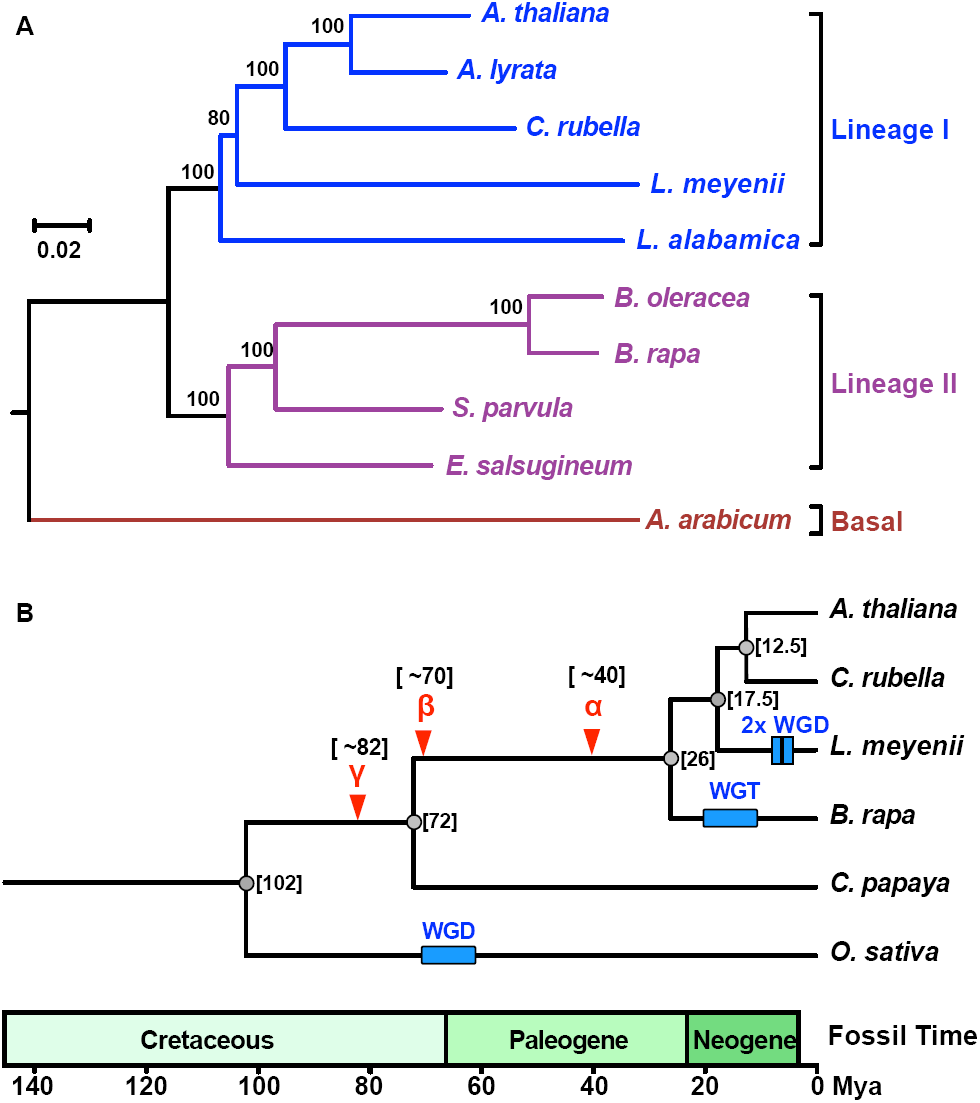
Phylogenetic analysis of representative members in different lineages of Brassicaceae family. (A) Phylogenetic tree and divergence time of six plant species. The tree was generated from 464 single orthologous genes using the maximum-likelihood method. Time (Mya) is listed in brackets, and corresponds to the time scale. Approximate positions of the paleo-WGD events (α, β, γ) are indicated by inverted triangles. Approximate positions of the species-specific WGD events are indicated by squares. (B) Representative members in different lineages of Brassicaceae family. The tree was generated from 464 single orthologous genes using the maximum-likelihood method. The scale bar shows expected number of amino acid substitutions per residue.

The Andes mountain range resides alongside the west coast of South America. In present day, the central part of the Andes comprises alpines, plateaus, highland lakes and valleys, which spans ∼400 kilometers wide with an average elevation of 4000 meters (Garzione et al. 2008). With regard to the formation of major high mountain ranges in the world, the process was usually steady and progressive over thousands and millions of years. But the Andes, especially the central Andes, uniquely experienced a spurt of elevation up to 3000 meters between 5 and 10 million years ago (Mya) (Garzione et al. 2008). This rapid uplift of the Andes profoundly shaped the climate and biodiversity of various ecosystems in the continent (Antonelli et al. 2009; Hoorn et al. 2010).

In the event of rapid environmental changes, plants could rely on the dispersal of propagules to establish new colonies in desirable habitats, or undertake phenotypic plasticity to mitigate disadvantageous environmental stimuli (Jump and Penuelas 2005; Chevin et al. 2010). However, the success of propagule dispersal and the magnitude of phenotypic plasticity both have limits (Jump and Penuelas 2005; Chevin et al. 2010). Given the existence of genetic variability and the heritability of genetic traits in a population, genetic evolution should be instrumental for plants to survive new local environmental conditions (Jump and Penuelas 2005; Chevin et al. 2010).

Polyploidy, which is a common phenomenon as a result of whole genome duplication (WGD), acts as a critical mechanism to encourage adaptation to harsh environment in plants (Chen 2007; Jiao et al. 2011). For example, WGD in the Brassicaceae provided ample genomic resource for functional innovation through evolution of duplicated genes and/or rewiring of genetic networks (De Smet and Van de Peer 2012; McGrath et al. 2014). The genomic analysis of *Arabidopsis thaliana* identified three major paleopolyploidy events (α, β and γ WGDs) having occurred before ∼40 Mya in the core Brassicaceae taxa (Kagale et al. 2014; Liu et al. 2014). But more importantly, it was the subsequent species-specific neo-/mesopolyploidy events that laid out the foundations for the species diversification of the Brassicaceae in the Neogene era (Kagale et al. 2014).

The genus *Lepidium* in the Brassicaceae family originated in the Mediterranean area as diploid species (Toledo et al. 1998). They were eventually dispersed throughout the world in the late Cenozoic period (Toledo et al. 1998). Compared with its counterparts, maca is a unique Andean *Lepidium* species with a disomic octoploid genome (2n = 8× = 64) (Quiros et al. 1996). This genomic feature, together with its preference to living at 4000 meters above sea level, makes maca an appropriate model to study both WGD events and the attendant high altitude adaptation. Here, we report the high-quality assembly and detailed analysis of the maca genome.

## RESULTS

### The high-quality genome assembly of maca

The sequencing of the whole maca genome on the Illumina HiSeq 2500 platform yielded 2.4 billion reads in paired-end libraries (Supplemental Table 1 and 2). A 743 Mb draft *de novo* assembly of the reads covered 98.93% of the estimated maca genome (751 Mb, Table 1, Supplemental Fig. 2-4, Supplemental Table 3). The contig and scaffold N50 sizes were 81 kb and 2.4 Mb respectively, which were among the longest in the published plant genomes based on short Illumina reads (Table 1, Supplemental Table 4). 2,636 scaffolds with sizes ranging from 2 kb to the maximum 8.8 Mb covered 86% of the assembled genome, to which over 97% of the clean reads could be mapped.

**Table 1.** Summary of *L. meyenii* genomic features

|  |  |
| --- | --- |
| **Estimated genome size (Mb)** | **751** |
| **Assembly statistics** |  |
| De novo assembly size (Mb) | 743 |
| Number of N50 contigs | 2,418 |
| N50 contig length (bp) | 81,780 |
| Number of N50 scaffolds | 92 |
| N50 scaffold length (Mb) | 2.4 |
| Number of N90 contigs | 122,974 |
| N90 contig length (bp) | 175 |
| Number of N90 scaffolds | 93,194 |
| N90 scaffold length (Mb) | 175 |
| Transposable elements content | 47.65% |
| **Gene annotation statistics** |  |
| Total number of protein-coding genes | 51,339 |
| Total exon number | 283,828 |
| Average exon number per gene | 5.53 |
| Average exon size (bp) | 200.72 |
| Total intron length (bp) | 46,265,182 |
| Total number of non-protein-coding genes | 34,846 |

Transposable elements (TEs) constitute 47.65% of the maca assembly (Table 1, Supplemental Table 5), which is the highest among all sequenced Brassicaceae species. Breakdown of the maca TE statistics showed higher proportion of retrotransposons than that of *A. thaliana* and *Brassica rapa* (Supplemental Fig. 5 and 6, Supplemental Table 6).

Gene annotation on the basis of homology and *ab initio* prediction methods identified 51,339 protein-coding genes (Table 1, Supplemental Fig. 7 and 8, Supplemental Table 7) and 34,846 non-protein-coding genes (miRNA, tRNA, rRNA and snRNA; Table 1, Supplemental Table 8). Out of all predicted protein-coding genes, about 81.53% could be classified into 3,861 protein families in the Swiss-Prot database, and about 99.92% had homologs in the TrEMBL database (Supplemental Table 9).

Comparative analysis of *A. thaliana*, *B. rapa*, *Carica papaya*, *Capsella rubella*, *O. sativa* and maca genes identified a total of 25,488 homologous gene families, in which 7,606 gene families were shared by all six species, and 980 gene families were maca-specific (Supplemental Fig. 9; Supplemental Table 10). Further comparisons of these species revealed 9,713 expanded gene families (366 KEGG enriched) in the maca genome (Supplemental Fig. 10; Supplemental Table 11). It is worth noting that many expanded genes and gene families were involved in abiotic stress response (i.e. WRKY, *bZIP*, *NAC*, *MYB* and *Homeobox* transcription factor gene families; Supplemental Fig. 11-15), hormone signaling pathway (i.e. brassinosteroid signaling pathway; Supplemental Fig. 16; Supplemental Table 12) and secondary metabolite biosynthesis (i.e. glucosinolate biosynthesis; Supplemental Fig. 17; Supplemental Table 13).

These data collectively build up to the annotated whole-genome sequence of maca (database available at www.herbal-genome.cn).

### Two close-spaced maca-specific WGDs occurred about 6.7 Mya

Maca is phylogenetically categorized into the lineage I of the widely diverse Brassicaceae (Fig. 1A) (Kagale et al. 2014). It diverged from the diploid *A. thaliana* about 17.5 Mya, and *B. rapa* of the lineage II about 26 Mya (Fig. 1B).

Here we analyzed the octoploidy of maca by aligning its assembled genome to the diploid *A. thaliana* genome with MCScanX. A surprising 1:4 projection ratio between *A. thaliana* and maca syntenic blocks showed the evidence of two maca-specific WGD events (Fig. 2A, Supplemental Fig. 18-21). This result was supported by the self-genome comparisons of maca, from which most paralogous gene clusters shared syntenic relationships with three other gene clusters (Supplemental Fig. 22). Additionally, the fourfold degenerate site transversion (4dTv) analysis of maca, *A. thaliana* and *B. rapa* confirmed the WGD events with 4dTv distance of 0.05 in maca and the whole genome triplication (WGT) event with 4dTv distance of 0.16 in *B. rapa* (Fig. 2B). The time of the two maca-specific WGDs was estimated at around 6.7 Mya using the method described elsewhere (Simillion et al. 2002), coinciding with the rapid elevation of Andes from 1000 m to 4000 m between 5 and 10 Mya (Fig. 2B and 2D; Supplemental Fig. 1B).

**Figure 2.**
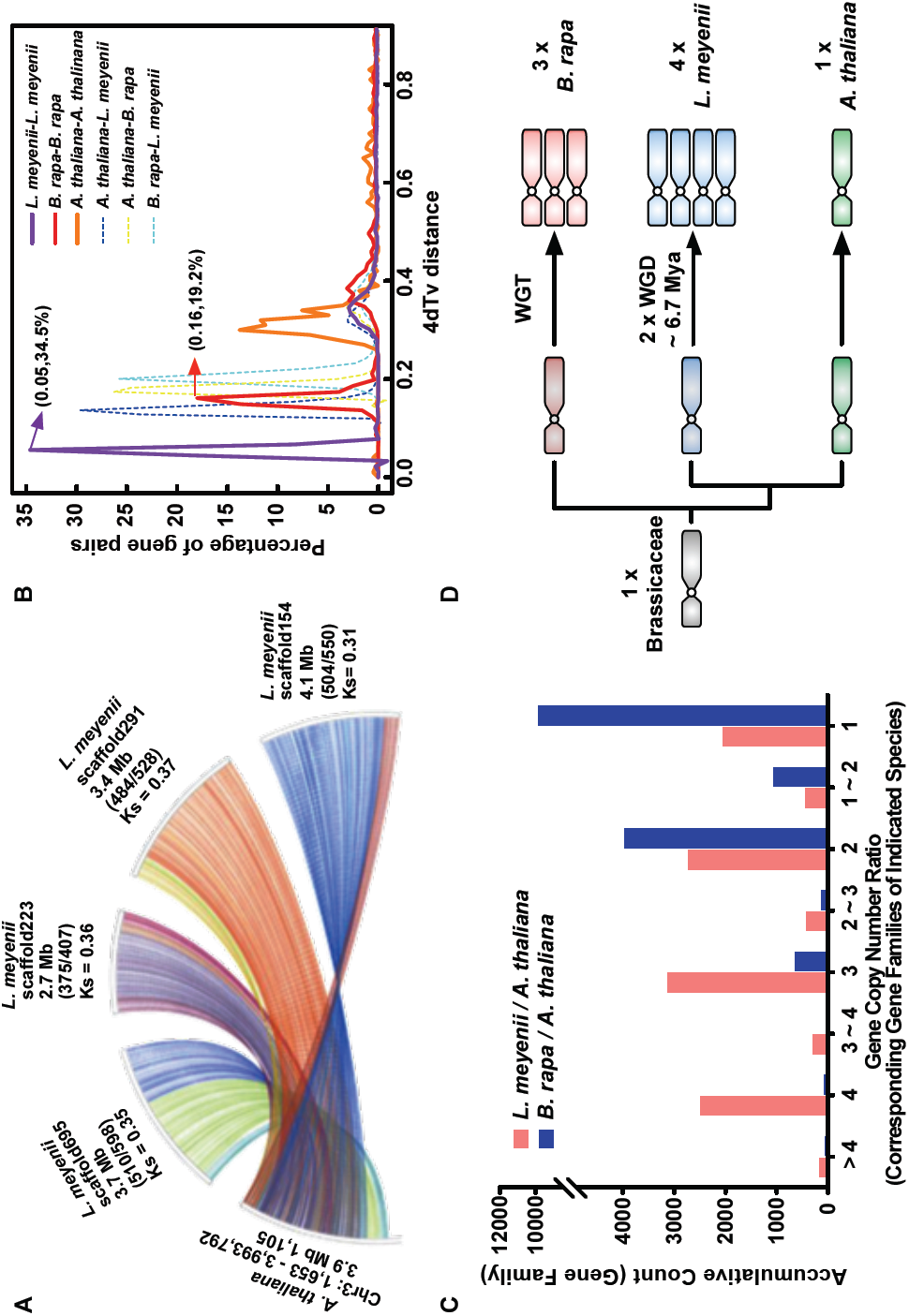
Whole genome duplications (WGDs) in the *Lepidium meyenii* genome at around 6.7 Mya. (A) *A. thaliana* chromosome 3 (1,653 – 3,993,792) shared syntenic relationships with four maca scaffolds (695, 223, 291 and 154, in different colors). Lengths of syntenic blocks, numbers of aligned genes out of total genes (in brackets) on the maca scaffolds, and the average synonymous rate (*Ks*) of all homologous gene pairs are shown. See Supplemental file for additional syntenic relationship graphs. (B) Fourfold degenerate site transversion (4dTv) analysis estimated that two maca-specific WGDs in the maca genome happened at around 6.7 Mya. (C) Gene copy number ratio of the corresponding gene families between the indicated species. (**D**) Schematic graph of genome evolution in *A. thaliana, L. meyenii* and *B. rapa.*

Comparisons of homologous gene numbers in the corresponding gene families of *A. thaliana, B. rapa* and maca genomes revealed higher degree of gene retention in maca after the two maca-specifc WGDs than that in *B. rapa* after WGT (Fig. 2C). This result suggests that the maca-specific WGDs provided ample genomic resource for the evolutionary adaptation to the harsh environment at high altitude Andes.

### Evolution of genes involved in leaf morphogenesis in the maca genome

An adaptive strategy for plants to cope with various abiotic stresses is to decrease leaf surface area through the modification of leaf morphogenesis (i.e. maca, Fig. 3A). During this intricate process, three major molecular mechanisms determine the leaf shape and complexity. On one hand, the class I *KNOTTED1-like homeobox (KNOX)* genes promote the leaflet initiation at the shoot apical meristem via influencing auxin-based patterning (Barkoulas et al. 2008; Blein et al. 2008; Hake and Ori 2002). In *A. thaliana*, the lobed-leaf morphology was manifest in transgenic clones overexpressing *KNOX* genes (Lincoln et al. 1994). The comparison of maca and *A. thaliana* genomes revealed ten homologous *KNOX* genes in maca as a result of the maca-specific WGDs (Fig. 3B), which provided genomic evidence for its lobate leaves. On the other hand, *CUP-SHAPED COTYLEDON (CUC)* and *REDUCED COMPLEXITY (RCO)* genes both regulate the formation of serrated leaf edges, but exert actions via distinct mechanisms. In *A. thaliana*, the evolutionary divergence of *CUC* gene gave rise to three paralogs (*CUC1*, *CUC2* and *CUC3*) as a result of neofunctionalization (Hasson et al. 2011). Despite their partially overlapping functions during leaf development, the balance between *CUC2* and its antagonist microRNA gene, *MIR164A*, primarily determines the degree of leaf edge serration (Nikovics et al. 2006). Tipping the balance toward *CUC2* leads to large and deep serration in *A. thaliana* (Larue et al. 2009). In the maca genome, the maca-specific WGDs yielded four homologous copies each for *CUC2* and *MIR164A* (Fig. 3C, Supplementary Fig. 23). Transcripts of all four maca *MIR164A* genes can be processed into similar *miR164a* (Supplementary Fig. 24). Interestingly, all four maca *CUC2* genes could not be silenced by *miR164a* due to the lack of functional target regions in the *CUC2* transcripts (Fig. 3C). This imbalance between *CUC2* and *MIR164A* may ultimately favor the serrated leaf edges in maca. Unaffected by the other two mechanisms, *RCO* genes, which appear in a cluster in the genome as a result of gene duplication from the *LATE MERISTEM IDENTITY 1 (LMI1)* type gene, sculpts developing leaflets to assist serration formation (Vlad et al. 2014). The simple leaf shape of *A. thaliana* results from the loss of *RCO* genes in the cluster (Fig. 3D) (Vlad et al. 2014). In the maca genome, *RCO* gene clusters were identified in the collinear regions of four scaffolds, indicating the retention of these genes after the maca-specific WGDs (Fig. 3D). Another *RCO* gene cluster was found in the non-collinear region of a fifth scaffold, which may be the result of a segmental duplication as evidenced by the MCScanX analysis (Fig. 3D). The abundance of *RCO* gene clusters, as well as the multiplication of *KNOX* and *CUC* genes may have collectively provided genomic explanation for the specialized leaf shape in maca.

**Figure 3.**
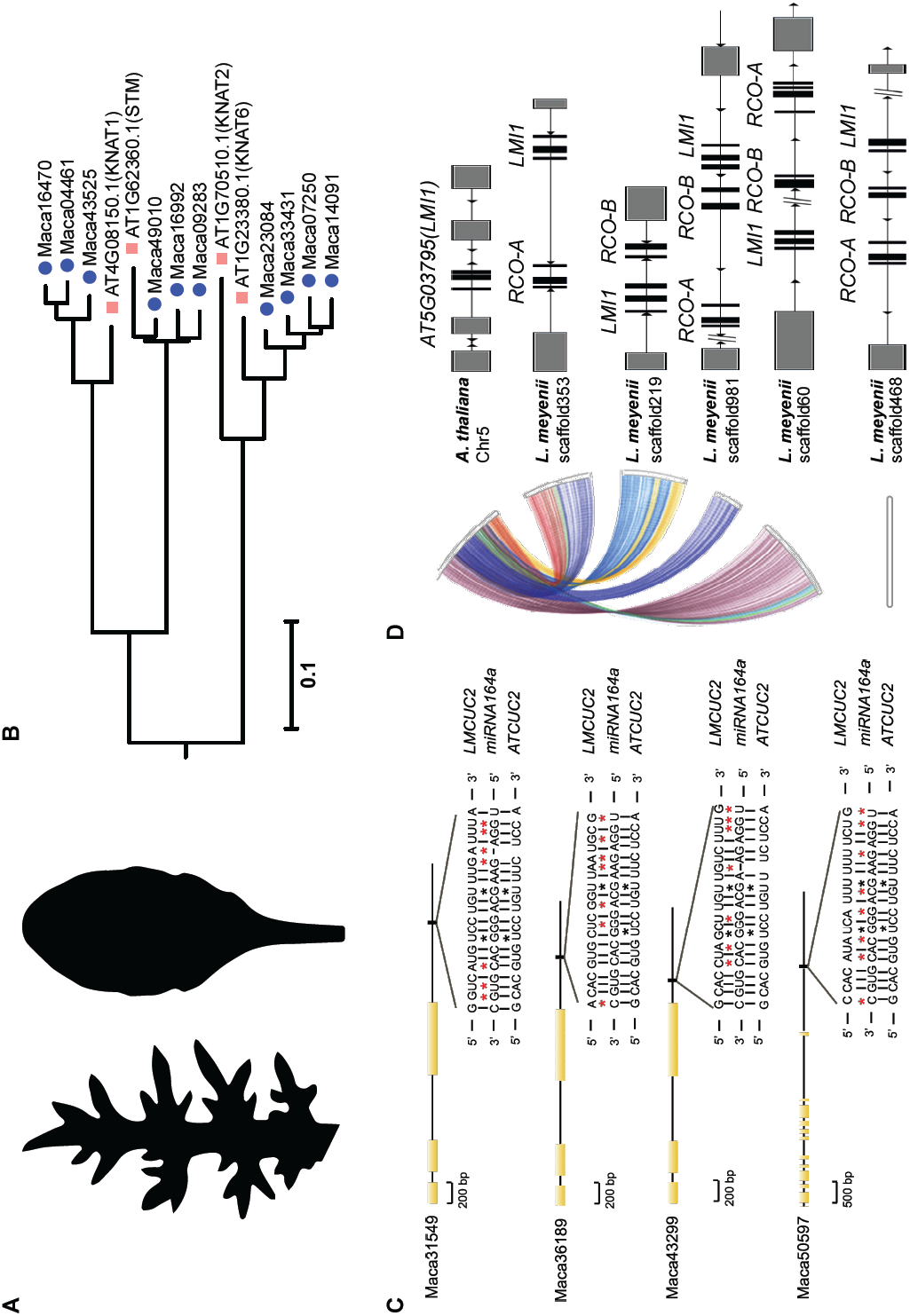
Evolution of genes involved in leaf morphogenesis in the maca genome. (A) Leaf silhouettes of maca (left) and *A. thaliana* (right). (B) Phylogenetic tree of class I *KNOTTED1-like homeobox* (*KNOX*) gene family from *L. meyenii* and *A. thaliana.* The scale bar shows expected number of amino acid substitutions per residue. (C) Four maca *CUC2* genes after two WGDs lost functional target sequences of *miR164a* in the *3’UTR.* Yellow boxes show exons of maca *CUC2* genes. The black box shows the position of the *miR164a* binding site, and the nucleotide sequence is detailed. Black asterisks represent the two mismatch sites allowed for the binding of *miR164a* to wild type *CUC2* transcript (*ATCUC2*). Red asterisks represent additional mismatches in maca *CUC2* transcript (*LMCUC2*) that disrupt *miR164a* binding. (D) Multiplication of *RCO/LMI1* gene clusters in the maca genome after two maca-specific WGDs. The *A. thaliana RCO/LMI1* gene cluster corresponds to the *RCO/LMI1* gene clusters in four maca syntenic blocks. A fifth gene cluster was identified in maca scaffold468. Black boxes represent exons in the genes.

### Evolution of genes involved in plant cold and UV-B adaptation pathways

As a perennial herbaceous plant growing on tropical highland, maca requires strong physiologic protection against frequent low nocturnal temperatures and long exposure to UV-B radiation. Despite the differences in adaptation strategies among plant species, it was proposed that: (1) most cold-adapted plants rely on a complicated cold acclimation process (Thomashow 1999) and/or a variety of antifreeze proteins (Preston and Sandve 2013; Griffith and Yaish 2004) to reduce freezing injuries in the event of low temperatures; (2) UV RESISTANCE LOCUS 8 (UVR8) protein-mediated photomorphogenic responses are essential for the establishment of UV-B tolerance in plants (Li et al. 2013). In view of the harsh growing conditions, maca is expected to retain key genes involved in these acclimation pathways after the maca-specific WGDs. Indeed, the examination of maca and *A. thaliana* genomes confirmed the multiplication and retention in the maca genome of antifreeze protein homologs (i.e. *BASIC CHITINASE* and *POLYGALACTURONASE INHIBITING PROTEIN 1*; Fig. 4), key genes in the well-characterized cold acclimation process (i.e. *CALMODULIN BINDING TRANSCRIPTIONAL ACTIVATORS*, *C-REPEAT BINDING FACTOR* and *INDUCER OF CBF EXPRESSION*; Fig. 4) and the UVR8-mediated signaling pathway (i.e. *UVR8*, *CONSTITUTIVELY PHOTOMORPHOGENIC 1*, *ELONGATED HYPOCOTYL 5* and *REPRESSORS OF UV-B PHOTOMORPHOGENESIS*; Fig. 4). These results implied the evolutionary role of the maca-specific WGDs in the development of acclimation traits in maca during Andes uplift.

**Figure 4.**
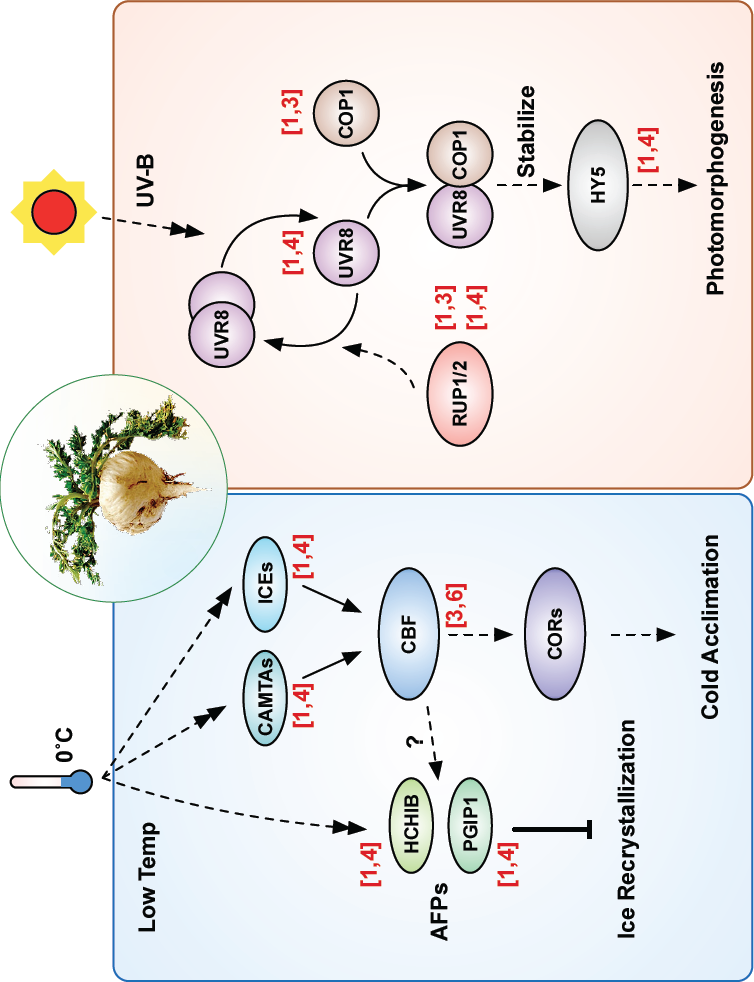
Evolution of genes involved in plant abiotic stress adaptation and flowering. The numbers of corresponding genes in *A. thaliana* and maca are shown in parentheses in the format of [*A. thaliana*, maca]. CAMTAs, CALMODULIN BINDING TRANSCRIPTIONAL ACTIVATORS; CBF, C-REPEAT BINDING FACTOR; ICE, INDUCER OF CBF EXPRESSION; CORs, COLD RESPONSIVE proteins; AFPs, antifreeze proteins; HCHIB, BASIC CHITINASE; PGIP1, POLYGALACTURONASE INHIBITING PROTEIN 1; UVR8, UV RESISTANCE LOCUS 8; COP1, CONSTITUTIVELY PHOTOMORPHOGENIC 1; HY5, ELONGATED HYPOCOTYL 5; RUP1/2, REPRESSOR OF UV-B PHOTOMORPHOGENESIS 1/2.

### Loss of important vernalization gene *FRIGIDA* in the maca genome

In order to successfully complete the life cycle, plants need to synchronize major reproductive events with two essential environmental cues – temperature and light (Franklin et al. 2014). For example, extended exposure to non-freezing cold temperatures prepares many winter plants for flowering in the upcoming spring (long day and warm temperature) (Amasino 2004). This *FRIGIDA (FRI)*-mediated process, known as vernalization, is apparent in winter-annual *A. thaliana* (Shindo et al. 2005). Loss of function mutation in the *FRI* gene is sufficient to disrupt vernalization and lead to rapid-cycle *A. thaliana* accessions (Johanson et al. 2000). Due to little seasonal change in average temperature and day lengths on tropical highland Andes, maca do not manifest vernalization process. In line with this phenomenon, the maca genome has lost all *FRI* homologous genes in the collinear regions compared with *A. thaliana* (Fig. 5), showing the loss of genes even after the maca-specific WGDs. Despite lacking functional *FRI* homologous genes, maca retains late flowering trait. This interesting phenomenon suggests that additional molecular mechanisms may play dominant roles in the regulation flowering time in maca.

**Figure 5.**
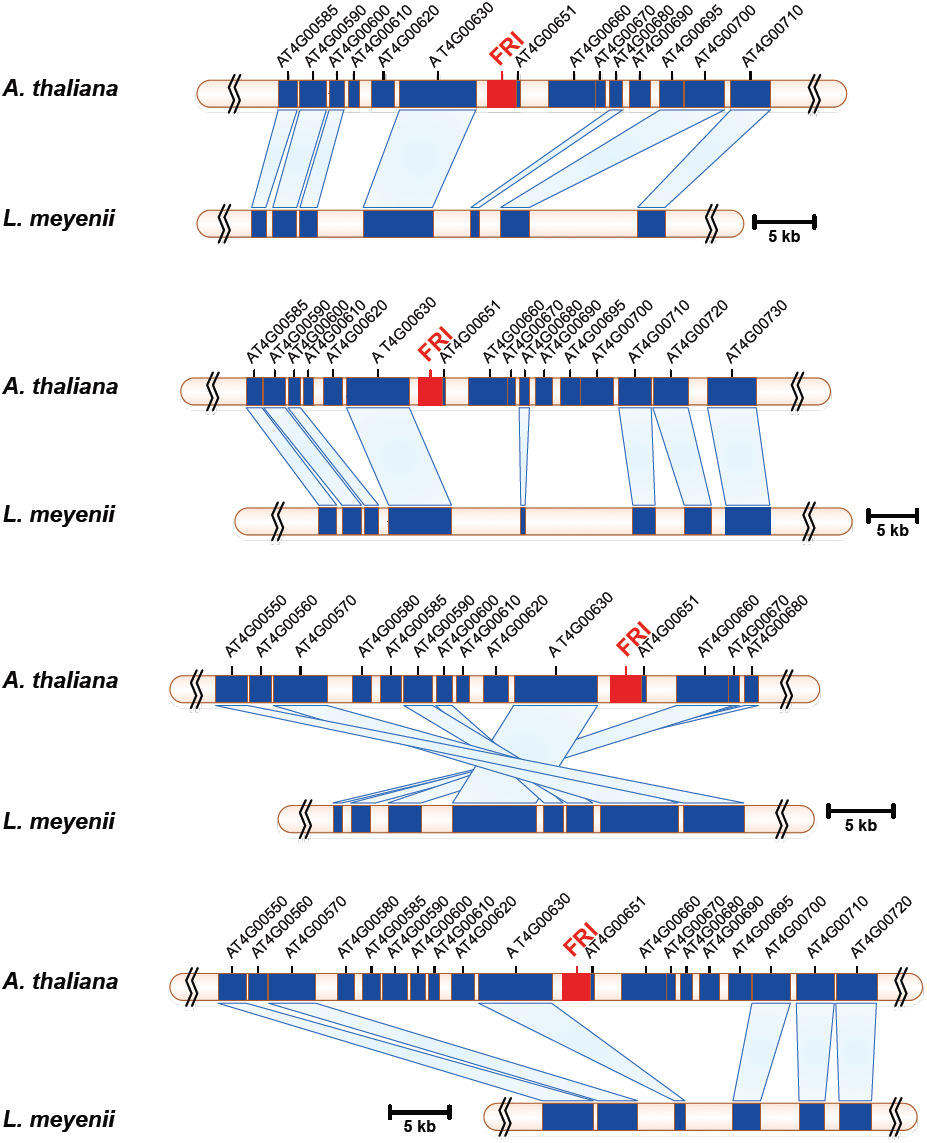
Zoomed-in views of *A. thaliana* and maca genome segments showed the losing of *FRI* genes in maca. Homologous gene pairs in *A. thaliana* and maca genomes are linked by light blue blocks.

### Genes under positive selection in maca

Increased rate of nonsynonymous substitution (*Ka*) relative to synonymous substitution (*Ks*) within certain genes may account for the adaptive evolution of organisms at the molecular level (Qiu et al. 2012). Comparisons of orthologous gene pairs identified 92 genes in maca versus *A. thaliana,* and 88 genes in maca versus *B. rapa*, having the ratio of *Ka/Ks* greater than 1.0 (*p*-value < 0.05, Supplemental Table 14 and 15). Interestingly, three genes with known physiological functions, including *MYB59*, *HISTONE DEACETYLASE 9* (*HDA9*) and *GLYCEROPHOSPHODIESTER PHOSPHODIESTERASE 5* (*GPDP5*), were present in both lists of genes under positive selection (Supplemental Fig. 25-27; Highlighted in Supplemental Table 14 and 15). In particular, *MYB59* encodes an R2R3-type transcription factor that regulates root development during the S phase of the cell cycle progression (Mu et al. 2009). Functional *HDA9* gene enhances seed dormancy and reduces seed germination rate (van Zanten et al. 2014). The product of *GDPD5* facilitates the release of inorganic phosphate from glycerophosphodiesters in the course of phosphate starvation (Borner et al. 2003). Rapid evolution of these three genes might have helped maca adapt to high altitude environment in terms of developing tuber root, seeding and enduring nutrient starvation.

## DISCUSSION

In this study, we have reported the high-quality assembly and detailed analyses of the disomic octoploid maca genome. By comparing the maca with the diploid *A. thaliana*, we identified two maca-specific WGD events in the maca genome, and estimated their occurrence at around 6.7 Mya during the uplift of the Andes. Subsequent analyses revealed that these two WGDs provided abundant genomic resources for reshaping maca traits and morphology in adaptation to the harsh environment. In this regard, the whole-genome sequencing of maca not only helps accelerate genetic improvement of important ergonomical traits for Brassicaceae crops, but also more importantly for the first time, provides a genomic basis for the high altitude adaptation in plants.

High altitude environment poses a variety of stress factors for living organisms, which include cold, strong UV-B radiation, low oxygen level and capricious climate. For human and highland animals, hypoxia and its related complications are the most severe physiological challenges (Cheviron and Brumfield 2011; Gou et al. 2014). Even though the ancient WGD events in vertebrates might have contributed to their current phenotypic complexities and evolutionary success (Freeling 2006), accumulating evidence showed that high altitude adaptation in human and animals mainly arose from rapid evolution of key genes and metabolic pathways for the oxygen uptake and delivery (Cheviron and Brumfield 2011; Qiu et al. 2012; Gou et al. 2014).

In contrast, widespread WGD events during the course of evolution played a major role in the species divergence and adaptation to new habitats in land plants (De Smet and Van de Peer 2012). For instance, the polyploid establishment in an allotment of monocots and eudicots coincided with the environmental disturbance around the Cretaceous-Paleogene extinction event, indicating that polyploidization may help mitigate the adverse effects of environmental stress in plant (Vanneste et al. 2014a; 2014b). Interestingly in the maca genome, two maca-specific WGDs occurred during the rapid uplift of the Andes. Since the genus *Lepidium* of Brassicaceae family originates in the Mediterranean area as diploid, this polyploidization in maca provides an excellent example on the enhancement of evolutionary fitness via WGD in the event of environmental change.

Polyploid plants are well known for the increased tolerance to abiotic stresses. In tetraploid *A. thaliana*, potentiated drought and salinity tolerance partly results from the alterations in cell proliferation, organ size and stoma physiology (del Pozo and Ramirez-Parra 2014; Chao et al. 2013). Despite these observations, genomic basis for the adaptive advantages after WGD remains unclear. One mathematic modeling of yeast cells showed that beneficial point mutations with strong fitness effects could accelerate the evolutionary adaptation of polyploid organisms (Selmecki et al. 2015). A second evolutionary model with yeast, however, concluded that duplication of a whole pathway was necessary for increased fitness (van Hoek and Hogeweg 2009). In fact, evidence supporting both hypotheses was obvious in the maca genome. On one hand, many maca genes under strong positive selection were involved in the development and abiotic stress response. On the other hand, expansion of cold and UV-B adaptation pathways, hormone signaling, and secondary metabolite biosynthesis pathways substantially strengthened the tolerance of maca to harsh environment. As the genomic resource multiplies after WGDs, the redundant pathways and networks are subject to massive gene loss, rewiring and reorganization for the emergence of new functions (De Smet and Van de Peer 2012). Indeed, multiplication and evolution of genes in leaf morphogenesis and vernalization leads to the specialized leaf shape and flowering time in maca. However, more research is warranted to observe these genes from the perspectives of pathways and networks.

In sum, findings from the maca genome confirmed the success of polyploidization of genomes in periods of ecological upheaval, and offered insight into the acquirement of adaptive advantage within short evolutionary period in Andean plants.

## METHODS

### Plant materials and DNA extraction

Maca plants were planted in the 4200m highland fields in Yunnan province of northwestern China. Three grams of fresh leaves from a 5-month-old individual were used to obtain genomic DNA. Genomic DNA was extracted with Qiagen DNeasy Plant Mini kits (Qiagen, Maryland, USA) following the manufacturer’s protocol. In total, about 100 μg high quality DNA were used for library construction.

### DNA library construction and sequencing

Genomic DNA was extracted and then fragmented randomly. After electrophoresis, DNA fragments of specific lengths were gel purified. In general, we constructed the libraries with insertion-sizes 200 bp, 308 bp (3 sets), 456 bp, 735 bp, 2 Kb, 5 Kb, 10 Kb and 20 Kb (Supplemental Table 1). Adapter ligation and DNA cluster preparation were performed and subjected to Illumina Genome HiSeq 2500™ sequencing with PE-100 protocol. We generated 321.54 gigabases (Gb) of data, which were equal to 428.72-fold coverage of the estimated 750 megabases (Mb) genome of *Lepidium meyenii* (Supplemental Fig. 2).

### Genome assembly

<u>Data processing.</u> For Illumina raw data, the processing of raw data included image analysis, base calling and sequence analysis. (1) Image analysis: Based on the TIF image files that came from the Illumina HiSeq 2500™ analyzer, we preliminarily detected the intensity of the cluster, got the position of each cluster and evaluated the background noise level; (2) Base calling: According to the evaluation results of cluster intensity and noise, we transformed the TIF image files into base sequence; (3) Sequence analysis: We removed substandard data according to the quality standards, and obtained sequence files and data quality assessment.

<u>Data filter.</u> The quality required is far higher for *de novo* assembly than for re-sequencing. In order to facilitate the assembly work, we took a series of checking and filtering measures on reads following the Illumina pipeline. We used the following criteria to filter low-quality reads:

1. Filter reads in which more than 5 percent of bases were N or poly A.
2. Filter low quality reads in which more than 30 bases were low quality.
3. Filter reads with adapter contamination.
4. Filter reads with small size.
5. Filter PCR duplicates. When read1 and read2 of two paired end reads were totally identical, these reads can be considered duplicates and were removed.

<u>Assembly.</u> The reads after the above filtering (result shown in Supplemental Table 2) and correction steps were used to perform assembly with SOAPec_v2.01 (http://soap.genomics.org.cn) developed by BGI. We used varies methods to assemble the maca genome. The best results were produced by SOAPdenovo. First, we tested a series of different *k-*mer (Supplemental Fig. 3) to construct contigs, and then picked the best results (*k*-mer = 87) to construct scaffolds. Another series of *k*-mer were used while scaffolding (Supplemental Fig. 4), and the best results (*k*-mer = 57) were used for the gapcloser process. We obtained a draft maca genome of 743 Mb, which is very close to the estimated size of the 750 Mb genome. The statistics for the final assembly was shown in Supplementary Table 4.

### Estimation of maca genome size with *k-* mer

*K*-mer refers to an artificial sequence division of *K* nucleotides. A raw sequencing read with L bp contains (L-K+1) *k*-mers if the length of each *k*-mer is *K* bp. The frequency of each *k*-mer can be calculated from the genome sequencing reads. The *k*-mer frequencies along the sequencing depth gradient follow a Poisson distribution for a given data set. During deduction, the genome size G = *K*_num/Peak_depth, where the *K*_num is the total number of *k*-mers, and Peak_depth is the expected value of *k*-mer depth. In this study, *K* = 17. All the short pair ends reads data were retained for the 17-mer analyses. The 17-mer frequency distribution derived from the sequencing reads is plotted in Supplemental Figure 2, with the peak of the 17-mer frequency distribution at about 38. With the total *k*-mer count being 28,539,182,510, the maca genome size can be estimated by the formula as 751.03 Mb. The statistics for the *k*-mer analysis was shown in Supplementary Table 3.

### Repeats annotation

Tandem repeats were identified across the genome with the help of the program Tandem Repeats Finder (Benson 1999). Transposable elements (TEs) in the genome were identified by a combination of homology-based and *de novo* approaches. For homology-based prediction, known repeats were identified using RepeatMasker and RepeatProteinMask against Repbase (Repbase Release 16.10;http://www.girinst.org/repbase/index.html).RepeatMasker (Tarailo-Graovac and Chen 2009) was applied for DNA-level identification using a custom library. At the protein level, RepeatProteinMask was used to perform a RMBLAST search against the TE protein database. For *de novo* prediction, RepeatModeler (http://repeatmasker.org/) (Tarailo-Graovac and Chen 2009) and LTR FINDER (Xu and Wang 2007) were used to identify *de novo* evolved repeats inferred from the assembled genome.

The distribution of detected TEs in the maca genome was shown in Supplemental Tables 5 and 6. The distributions of the percentage of genome composed of TEs of differing divergence from consensus sequence (estimated from Repbase (Jurka et al. 2005) or TE library consensus) identified by the homology-based and the consensus sequence methods were shown in Supplemental Figures 5 and 6. The peak divergence rate of TEs was measured by the percentage of substitutions in well-aligned regions between annotated repeats in maca and consensus sequences in Repbase.

### Protein-coding genes prediction

To predict protein-coding genes in the maca genome, we used both homology-based and *de novo* prediction methods.

#### 1. Homology based gene prediction

Proteins of *Oryza sativa, Populus trichocarpa, Arabidopsis lyrata, Arabidopsis thaliana*, *Brassica rapa*, and *Eutrema salsugineum* were used in the homology-based annotation. All these proteins were obtained from Phytozome v9.1 (http://www.phytozome.net/). First, TBLASTN was processed with parameters of ‘*E*-value = 1e^-5^, -F F’. BLAST hits that correspond to reference proteins were concatenated by Solar software, and low-quality records were filtered. Genomic sequence of each reference protein was extended upstream and downstream by 2000 bp to represent a protein-coding region. GeneWise software (Birney and Durbin 2000) was used to predict gene structure contained in each protein region.

#### 2. *De novo* gene prediction

We used two *de novo* prediction programs: AUGUSTUS (Stanke et al. 2006) and GlimmerHMM (Majoros et al. 2004), with gene model parameters trained from *A. thaliana*. We filtered partial genes and small genes with less than 150 bp coding lengths.

#### 3. High-confidence gene model generation

The homology-based and *de novo* gene sets were merged to form a comprehensive and non-redundant reference gene set using the GLEAN software, which is a tool for creating consensus gene lists (Supplemental Fig. 7) by integrating gene evidence through latent class analysis.

Based on the above-mentioned methods, we obtained a maca gene set that included 51,339 protein-coding genes (Supplemental Table 7).

### Non-protein-coding genes annotation

The tRNAscan-SE (version 1.23) (Lowe and Eddy 1997) software with default parameters for eukaryote was used for tRNA annotation. rRNA annotation was based on homology information of plant rRNAs using BLASTN with parameters of ‘*E*-value = 1e^-5^’. The miRNA and snRNA genes were predicted by INFERNAL software (http://infernal.janelia.org, version 0.81) (Nawrocki et al. 2009) against the Rfam database (Release 11.0) (Gardner et al. 2011). The distribution of ncRNAs in the maca genome is shown in Supplemental Table 8.

### Gene function annotation

Gene functions were assigned according to the best match alignment using BLASTP against Swiss-Prot or TrEMBL databases (Bairoch and Apweiler 2000). Gene motifs and domains were determined using InterProScan (Mulder and Apweiler 2007) against protein databases including ProDom, PRINTS, Pfam, SMART, PANTHER and PROSITE. Gene Ontology IDs were obtained from the corresponding Swiss-Prot and TrEMBL entries. All genes were aligned against the KEGG proteins. The pathway to which the gene might belong was derived from the matching genes in KEGG. In total, 99.92% of the predicted genes were assigned with a functional annotation (Supplementary Table 9).

### Gene family cluster

Protein sequences for all protein-coding genes in *A. thaliana, B. rapa, C. papaya, C. rubella, O. sativa* were downloaded from Phytozome v9.1 (http://www.phytozome.net/). To identify gene family clusters in these species, all-versus-all protein searches were performed using BLASTP with the parameter of ‘*E*-value = 1e^-5^’. OrthoMCL package (Version 1.4) (Li et al. 2003) was used to process high-scoring segment pairs (HSPs). MCL software in OrthoMCL was used to define final paralogous and orthologous genes with the parameter of ‘-abc – I = 1.5’.

### Phylogenetic tree construction

Single copy genes (i.e, one copy in all species) were used to construct phylogenetic tree of *A. thaliana, B. rapa, C. papaya, C. rubella, O. sativa* and *L. meyenii*. Multiple sequence alignments were performed using MUSCLE (Edgar 2004). Fourfold degenerate sites were extracted from each gene and concatenated into a supergene for each species. MrBayes software (http://mrbayes.sourceforge.net, version 3.1.2) (Huelsenbeck and Ronquist 2001) was used to reconstruct phylogenetic trees between species.

### Divergence time estimation

The program MCMCTREE program, implemented in PAML package (Yang 2007), was used to estimate divergence time of *L. meyenii, C. rubella, A. thaliana, B. rapa, C. papaya,* and *O. sativa* (as an outgroup). The HKY85 model (model = 4) and independent rates molecular clock (clock = 2) were used for calculation. The MCMC process of MCMCTREE was performed with the samples 1,000,000 times, with a sample frequency setting of 2, after a burnin of 200,000. The fine-tune parameters in the control file were set to ‘0.0058 0.0126 0.3 0.43455 0.7’ to make the acceptance proportions fall into the interval (0.2, 0.4). The divergence age was set to 102 Mya, which was the corrected divergence time between *C. papaya* and *O. sativa* from the time tree database (http://www.timetree.org/). See Figure 1A.

### Expansion and contraction of gene families

Based on the calculated phylogeny and the divergence time, CAFÉ (Computational Analysis of Gene Family Evolution, version 2.1) (De Bie et al. 2006), a tool based on the stochastic birth and death model for the statistical analysis of the evolution of gene family size, was applied to identify gene families that had undergone expansion and/or contraction in the *L. meyenii, A. thaliana, B. rapa, C. papaya, C. rubella,* and *O. sativa* with the parameters ‘*p*-value = 0.05, number of threads = 10, number of random = 1000, and search for lambda’.

### Detection of positively selected genes

To detect genes under positive selection, we used the coding DNA sequence (CDS) libraries of *A. thaliana* and *B. rapa* to run BLASTN against the maca CDS library, respectively. The best hits with >80% identity and >80% alignment rate were run under LASTZ (http://www.bx.psu.edu/miller_lab/) with default parameters. The coding sequences of orthologs were aligned using MUSCLE software (http://www.drive5.com/muscle, version 3.7) with default parameters. The gene pairs were then analyzed in *KaKs*_Calculator (version 1.2) (Zhang et al. 2006) with default parameters.

### Detection of segmental duplication

Whole-genome comparisons were used to identify segmental duplications (SDs). Self-alignment of each genome assembly was performed using LASTZ (http://www.bx.psu.edu/miller_lab/) with the parameters set at ‘T = 2, C = 2, H = 2000, Y = 3400, L = 6000, K = 2200’. Before alignment, scaffolds shorter than 1000bp were discarded. After alignment, repeat sequences in syntenic blocks with > 500 bp length and > 85% identity were unmasked. LASTZ analysis was performed again with the above-mentioned parameters. Two sequences (query and target) with > 1kb length and 90% ∼98% identity were defined as SDs.

## DATA ACCESSION CODES

The assembly and annotation of the maca genome are available at http://www.herbal-genome.cn. The sequencing reads of each Illumina sequencing libraries have been deposited under GenBank with Project ID PRJNA264949. All short-read data are available via Sequence Read Archive: SRR1631197, SRR1631211, SRR1631213, SRR1631216, SRR1631217, SRR1631219, SRR1631222, SRR1631223, SRR1631224, SRR1631225.

## ACKNOWLEDGEMENT

This work was supported by the Pilot Project for Establishing Social Service System by Agricultural Science and Education in Yunnan Province, Medical Plant Unit (2014NG003).

## AUTHOR CONTRIBUTIONS

J.Z., Y.D., W.C., J.S. and W.W. planned and coordinated the project and wrote the manuscript. Y.T., S.H., X.M. and L.Y. collected and grew the plant material. J.Z., Y.D and G.Z. sequenced, processed the raw data, assembled and annotated the maca genome. X.W. and Y.Z. analyzed the genomes. J.J.Z. set up the database. Y.T., N.L. J.Z., Y.W. and Y.M conducted genome evolution analysis. Y.X. gave helpful suggestion on the whole project. J.Z., S.Y. and G.Z analyzed the genes/gene families related to specific traits. L.Z. conducted whole genome duplication related analysis.

